# Open-Source Framework for Measurement of the Gradient Impulse Response Function

**DOI:** 10.64898/2026.08.24.745510

**Authors:** James B. Bacon, Rudy Rizzo, Simon M. Finney, C. John Evans, Fabrizio Fasano, Peter Jezzard, William T. Clarke

## Abstract

The Gradient Impulse Response Function (GIRF) is widely used to model and correct gradient system imperfections in MRI, but scanner-specific GIRF measurement remains inaccessible to many research groups because existing approaches rely on specialised field monitoring hardware or fragmented and non-reproducible software workflows. To address this limitation, an open-source, end-to-end framework for phantom-based GIRF measurement is presented, providing a reproducible workflow requiring only standard MRI hardware and a spherical water phantom. The framework integrates vendor-independent pulse sequence generation, phantom-based data acquisition, automated data processing, and GIRF estimation. The framework was validated by comparing GIRF-predicted non-Cartesian k-space trajectories with independent measurements acquired using NMR field probes, which served as the gold-standard for trajectory characterization. Accurate prediction of rosette and spiral trajectories was demonstrated across multiple imaging orientations, with substantially lower trajectory error than the corresponding nominal trajectories. By providing the first openly available end-to-end implementation for phantom-based GIRF measurement, the barrier to routine scanner-specific GIRF characterisation is reduced, facilitating broader adoption of GIRF-based methods across the MRI community.

## Introduction

Non-ideal gradient system behaviour, which includes eddy currents, mechanical resonances, and timing delays, causes the gradient fields generated during an MR experiment to deviate from those intended by the prescribed gradient waveforms [1]. These deviations degrade image quality, giving rise to artefacts such as geometric distortion, blurring, and ghosting [2]. Their impact is particularly pronounced in non-Cartesian imaging, where high-duty-cycle, rapidly switching gradient waveforms make image reconstruction highly sensitive to errors in the realised k-space trajectory (the time-integral of the gradient waveforms) [3]. In addition to non-resonant eddy-current effects, mechanical vibrations of the gradient system introduce frequency-dependent oscillations in the generated fields that are not fully compensated by conventional vendor-implemented pre-emphasis schemes, which typically model eddy currents solely as a sum of exponential decays [4,5]. Accurate characterisation of the dynamic response of MRI gradient systems is therefore essential for predicting and correcting gradient-induced field perturbations and their associated image artefacts.

The Gradient Impulse Response Function (GIRF) provides a linear and time invariant systems model of the dynamic response of an MRI gradient system [6]. Under the assumption that the dynamic response is linear and time-invariant [7,8], the GIRF characterises how prescribed gradient waveforms are transformed into the realised gradient and B_0_ fields, enabling prediction of gradient-induced field perturbations. Consequently, the GIRF has been widely adopted to predict and compensate for the effects of non-ideal gradient system behaviour, including prediction of k-space trajectory deviations for non-Cartesian image reconstruction [9,10], improvement of gradient pre-emphasis [11–14], optimisation of multiband pulse design [15], and compensation of frequency-dependent field oscillations in MR spectroscopy [16]. These diverse applications have established the GIRF as a versatile tool for modelling and correcting gradient system imperfections.

Despite the widespread adoption of the GIRF, obtaining scanner-specific GIRF measurements remains a significant practical challenge. Existing approaches often rely on specialised field monitoring hardware [6], which is not routinely available, or provide only individual components of the measurement workflow, such as GIRF estimation routines without the accompanying pulse sequences [11,17]. Although phantom-based measurements have been described in the literature [3,18], there is currently no openly available framework that provides an integrated end-to-end workflow encompassing pulse sequence generation, phantom-based data acquisition, data processing, and GIRF estimation. Consequently, implementing a complete GIRF measurement workflow often requires considerable laboratory-specific software development, limiting accessibility and reproducibility.

In this work, we present an open-source framework for end-to-end phantom-based GIRF measurement. By integrating pulse sequence generation, data acquisition, and GIRF estimation into a single reproducible workflow, the framework lowers the barrier to routine scanner-specific GIRF characterisation using standard MRI hardware. The resulting GIRFs are validated through comparison of GIRF-predicted k-space trajectories with independent measurements acquired using NMR field probes [19].

## Methods

The proposed workflow is illustrated in Figure 1. The framework comprises three principal components: phantom-based acquisition of gradient response measurements, automated data processing, and GIRF estimation. Following generation of a vendor-independent Pulseq sequence [20], measurements are acquired using a thin-slice method with 2D phase encoding [3,18]. The acquired data are subsequently processed through a series of automated reconstruction and phase-processing steps to estimate the corresponding gradient and B_0_ field responses, from which the GIRF is estimated in the frequency domain.

**Figure 1:**
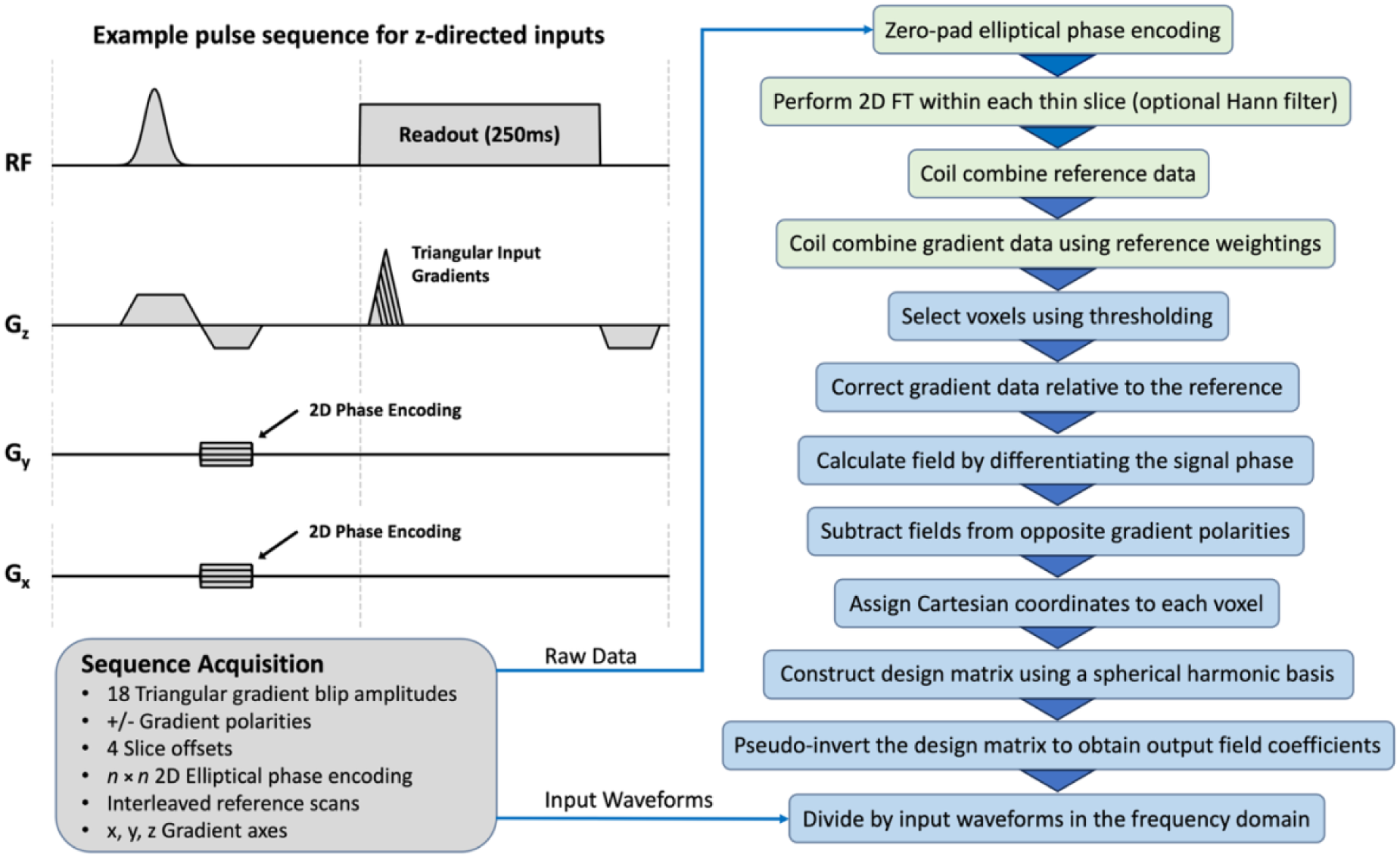
Overview of the proposed end-to-end framework for phantom-based GIRF measurement. A vendor-independent sequence is generated using PyPulseq *[21]* and used to acquire thin-slice, phase-encoded measurements from a spherical phantom. The acquired data are processed through an automated reconstruction pipeline to estimate the output field responses, which are subsequently transformed into the frequency domain to calculate the GIRF. The workflow is divided into three principal stages: pulse sequence generation (grey), automated data processing (light green), and GIRF estimation (blue).

The acquisition stage implements the phantom-based thin-slice method proposed by Duyn et al. [3] to characterise the gradient system response. Four parallel thin slices are excited at offset positions within a spherical phantom and encoded using 2D phase encoding [18], allowing the gradient-induced phase to be measured independently for each gradient axis (Figure 2). Following slice excitation, 18 triangular gradient blips are applied, each with both positive and negative polarity, after a delay of 2 ms. The gradient amplitudes are uniformly distributed from 9 to 39.6 mT/m with a fixed slew rate of 180 T/m/s [17]. Triangular blips were selected in preference to chirp-based excitations [22], as they provide greater spectral density at low frequencies. This improves estimation of the GIRF over the frequency range most relevant to MR spectroscopy, the application targeted by the authors, where gradient-induced field perturbations overlap with the metabolite resonance frequencies [16,23].

**Figure 2:**
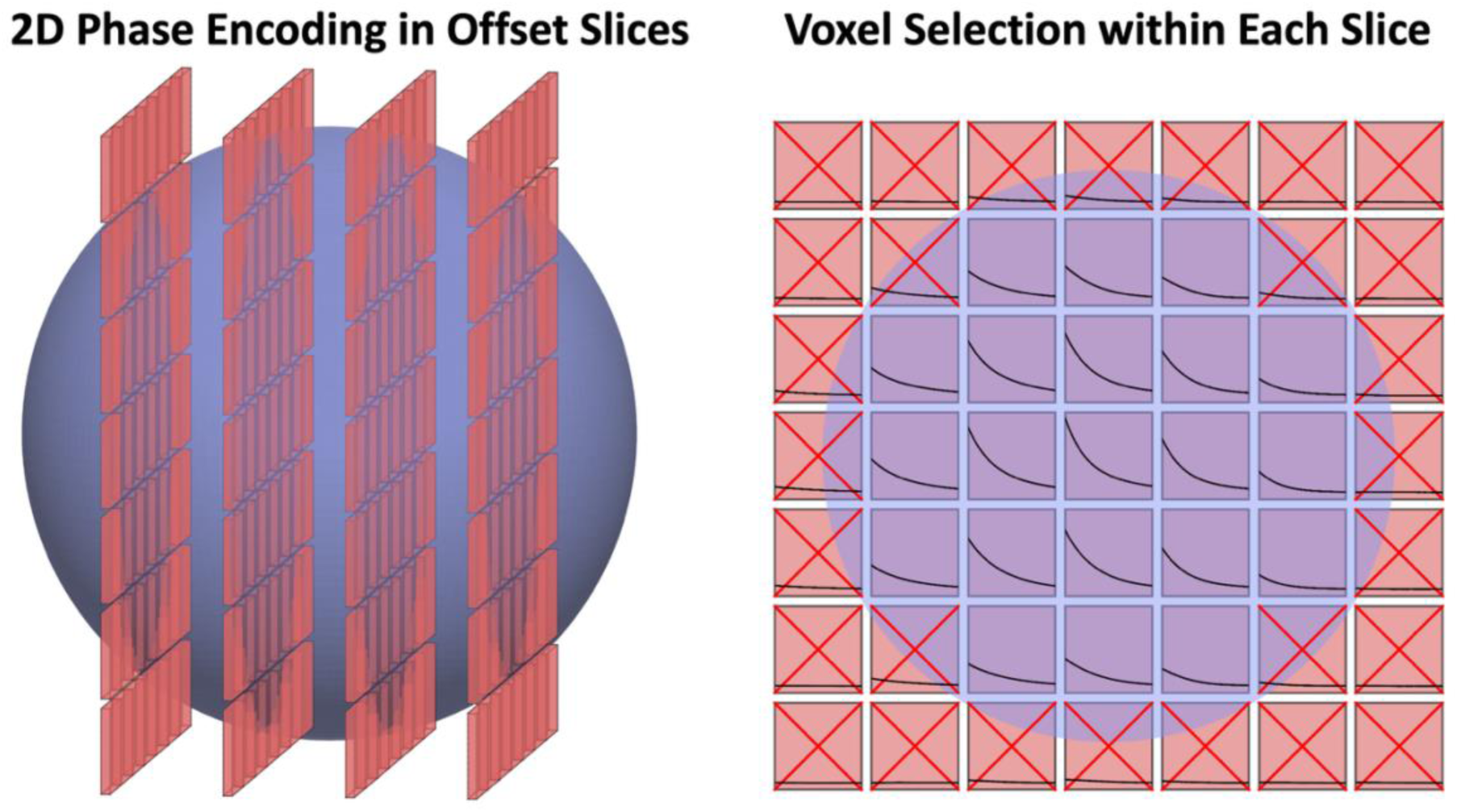
Thin-slice acquisition and voxel selection used for phantom-based GIRF measurement. Left: Four offset, phase-encoded thin slices provide spatial sampling of the gradient-induced field response throughout the spherical phantom. Right: Voxels are selected using an intensity threshold prior to spherical harmonic fitting. The threshold preferentially excludes the low-signal voxels near the phantom boundary. Figure adapted from Rahmer et al. *[18]*

Optional elliptical 2D phase encoding is used within each thin slice to spatially resolve the measured field response, enabling estimation of higher-order spherical harmonic components of the GIRF [18]. The reduced effective voxel size also minimises intravoxel dephasing, lengthening the T2* to permit acquisition windows of up to 250 ms, and thereby improving the GIRF spectral resolution. Measurements from the four offset slices are interleaved, resulting in an effective TR of 4.6 s per slice. Reference acquisitions without applied gradient blips are inserted every three gradient amplitudes to correct for residual phase contributions which can arise from the slice-selective and phase encoding gradients, as well as temporal system drift [17]. Using the default acquisition protocol with a 7 × 7 phase-encoding matrix, the total acquisition time is approximately 90 min per gradient axis. When only the zeroth-order (B_0_) cross-terms and self-response terms are required, the in-plane encoding can be omitted, reducing the acquisition time to approximately 3 min per axis.

Data processing and GIRF estimation follow the methodology described by Rahmer et al. [18], as summarised in Figure 1. Voxels with insufficient signal are excluded using user-guided intensity thresholding (Figure 2). For each retained voxel, the temporal phase evolution is differentiated to estimate the output field, while measurements acquired with opposite gradient polarities are subtracted to remove concomitant field effects and susceptibility-induced off-resonance contributions [24]. The measured fields are subsequently fitted to a spherical harmonic basis using their known voxel coordinates, and the resulting coefficients are used to estimate the GIRF in the frequency domain by dividing the output field components by the corresponding input gradient waveforms. The framework also supports optional truncation of the readout to optimise the trade-off between spectral resolution and noise amplification for different applications.

The framework is implemented entirely in Python and released as open-source software to promote accessibility and reproducibility. Vendor-independent pulse sequence generation is achieved using PyPulseq [21], while the remaining workflow is implemented as a modular processing pipeline comprising sequence generation, data processing, and GIRF estimation. The software is distributed with managed dependencies and accompanying documentation through a public GitHub repository (https://github.com/jbbacon/GIRF_PE_Python) to facilitate reproducible deployment.

The framework was validated by performing GIRF measurements on a research only 3T MAGNETOM Connectom Scanner (Siemens Healthineers, Forchheim, Germany). Measurements were acquired using an 11 cm diameter spherical HDPE phantom (Spherical Phantom Application Research Kit (SPARK), Gold Standard Phantoms, Sheffield, UK) filled with NiCl_2_-doped saline (deionized H_2_O, NiCl_2_: 5.5 mM, NaCl: 30.0 mM) [25]. The reduced phantom diameter was selected to minimize B_1_^+^ inhomogeneity. Although the SPARK phantom was used throughout this work, the framework is compatible with any spherical water phantom exhibiting a single resonance peak [16]. Measurements were acquired using slice offsets of ±8 and ±16 mm (1 mm slice thickness), a 7×7 elliptical phase encoding matrix, a 5 μs dwell time, and an 85° excitation flip angle.

To independently validate the estimated GIRFs, predicted k-space trajectories were compared with direct field measurements acquired using a NeuroCam 3T head coil incorporating field probe sensors (Skope Magnetic Resonance Technologies, Zurich, Switzerland), widely regarded as the gold-standard reference for direct and image-concurrent trajectory measurement. Two non-Cartesian MRSI trajectories were selected for validation: a rosette trajectory and a spiral trajectory [26]. Both trajectories utilized golden angle sampling between each petal/spiral (trajectory details given in [27]) and each trajectory was acquired in axial, sagittal and coronal orientations. These trajectories were selected because of their demanding gradient waveforms and long readout durations, making them particularly sensitive to gradient system imperfections. Predicted trajectories were generated by convolving the prescribed gradient waveforms with the estimated GIRFs [9]. Agreement was assessed qualitatively through trajectory overlays and quantitatively using the root-mean-square (RMS) trajectory error relative to both the nominal and measured trajectories throughout the readout.

## Results

The magnitudes of the GIRFs estimated using the proposed framework are shown in Figure 3. The GIRF is denoted by *H_j_*_,*m*_(*ω*), where *j* denotes the input gradient axis and *m* denotes the spherical harmonic component of the output field. Figure 3 displays the zeroth- and first-order spherical harmonic components estimated using a reconstructed readout duration of 70 ms, comparable to the acquisition windows typically achieved using NMR field camera measurements at 3T and under typical phantom shim conditions [6]. The corresponding phase of the self-response terms is provided in Figure S1.

**Figure 3:**
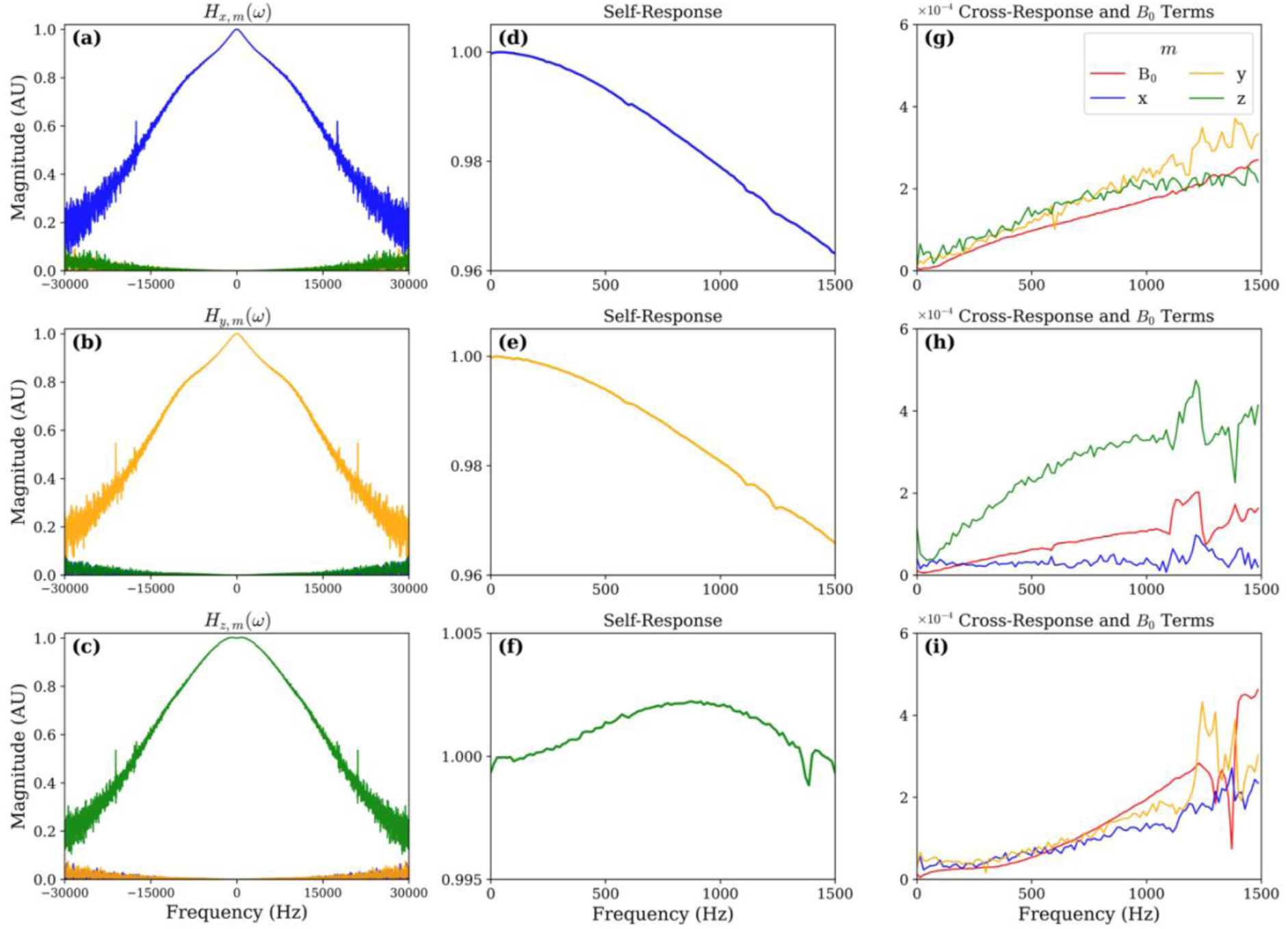
Magnitude of the GIRFs estimated on a 3T Connectom. **(a-c)** Full frequency response of the zeroth- and first-order spherical harmonic components for input gradient axes *j* = *x*, *y* and *z*, respectively. **(d-i)** Enlarged views of the corresponding self-response **(d-f)** and cross-response/*B*_0_ terms **(g-i)**, each displayed between 0 and 1500 Hz. All GIRFs were estimated using a reconstructed readout duration of 70 ms. Output field spherical harmonic components (*m*) are colour coded as *B*_0_ (red), *x* (blue), *y* (orange), and *z* (green).

For trajectory prediction, GIRFs estimated using reconstructed readout durations of 175 ms were used for the B_0_ and self-response terms, while, due to their lower sensitivity, the first-order cross-response terms were estimated using the first 100 ms of data only [16]. This trades reduced GIRF frequency resolution for higher SNR. Prior to trajectory prediction, a dwell-time compensation filter was applied to the estimated GIRFs to account for the finite sampling interval of the GIRF acquisition [19]. The compensated GIRFs were convolved with the prescribed gradient waveforms to predict the corresponding gradient outputs, which were subsequently integrated to obtain the predicted k-space trajectories. Figure 4 compares the nominal, GIRF-predicted and field probes measured trajectories for the rosette acquisitions in the axial plane. Corresponding results for the spiral trajectory, together with the coronal and sagittal orientations, are provided in Figure S2 and S3.

**Figure 4:**
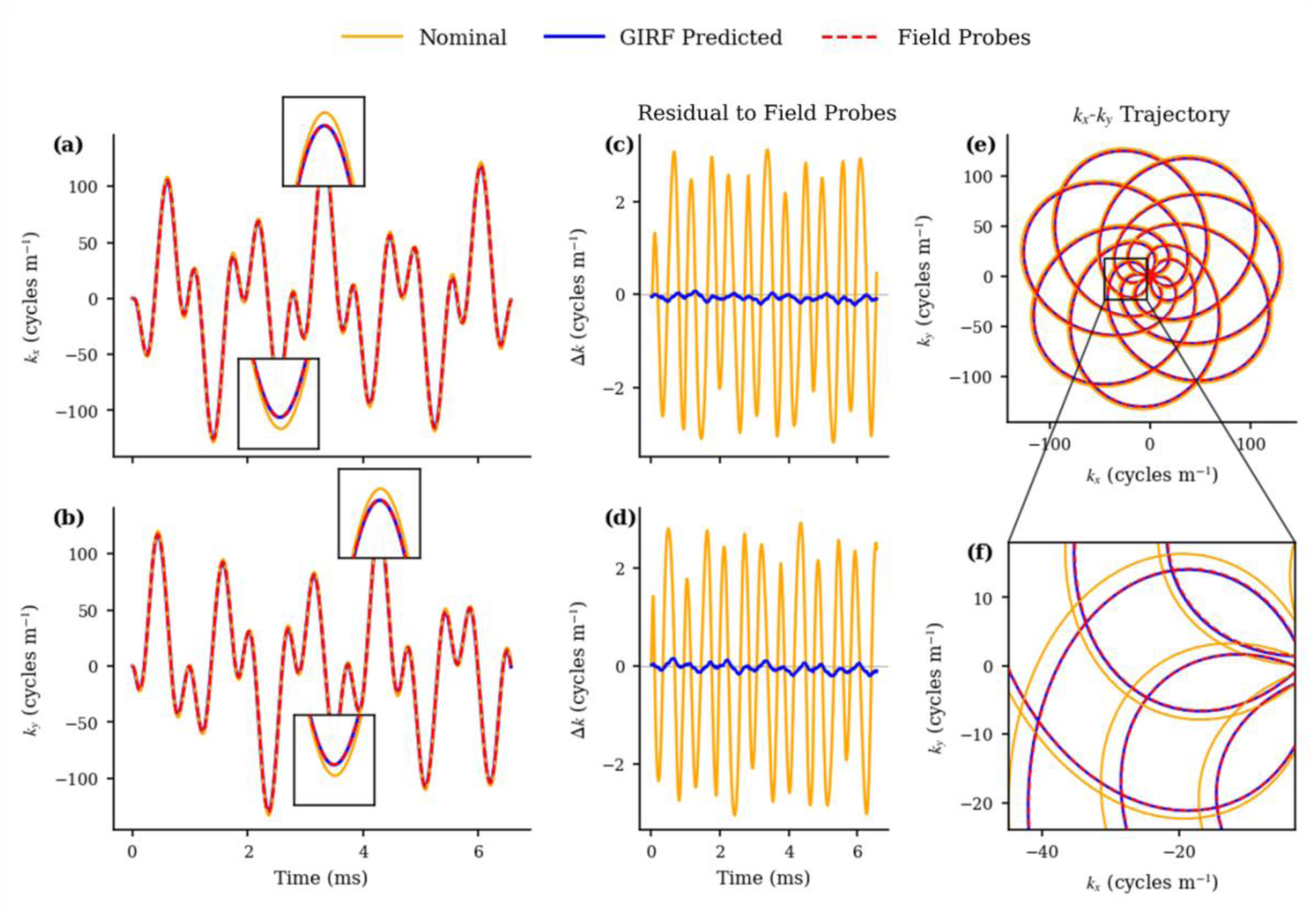
Comparison of the nominal, GIRF-predicted and field probes-measured rosette trajectories acquired in the axial plane. Only the first 6.6 ms of the readout is displayed. **(a,b)** *k_x_* and *k_y_* trajectories as a function of time, with enlarged views highlighting the agreement between the GIRF-predicted and measured trajectories. **(c,d)** Residual trajectory error relative to the field probes measurement for the nominal and GIRF-predicted trajectories. **(e)** Two dimensional *k_x_*-*k_y_* trajectory. **(f)** Enlarged view around the central region of the trajectory. Trajectories are colour coded as Nominal (orange), GIRF-predicted (blue), and Field Probes measured (red).

Figure 4 demonstrates close agreement between the GIRF-predicted and field probes-measured trajectories throughout the first 6.6 ms of the readout. Relative to the nominal trajectory, the GIRF prediction substantially reduced the trajectory error, as reflected by both the temporal trajectory profiles and the residual error. Quantitatively, the RMS trajectory error was reduced from 2.62 cycles m^-1^ to 0.13 cycles m^-1^ when using the GIRF-predicted trajectory, representing a 95% reduction. Similar improvements were observed for the spiral trajectory and remaining imaging orientations, with the mean RMS trajectory error across all trajectories and orientations decreasing from 2.78 cycles m^-1^ to 0.18 cycles m^-1^ over this readout duration using the GIRF prediction.

To assess the agreement between the predicted and measured trajectories over the conventional acquisition window supported by NMR field probes, the comparison was extended to a 90 ms readout (Figure 5a). Throughout the acquisition, the GIRF-predicted trajectories remained in closer agreement with the field probes measurements than the nominal trajectories. However, over the longer readout there was a systematic linear trend in the trajectory error. For both the *k_x_* and *k_y_* trajectories, the nominal and GIRF-predicted residuals shared a common linear trend despite the substantially smaller residual magnitude obtained using the GIRF prediction. Over the full 90 ms acquisition, the corresponding RMS trajectory error increased to 3.07 cycles m^-1^ for the nominal trajectory and 1.45 cycles m^-1^ for the GIRF-predicted trajectory. This behaviour was observed consistently across both trajectory types and all imaging orientations.

**Figure 5:**
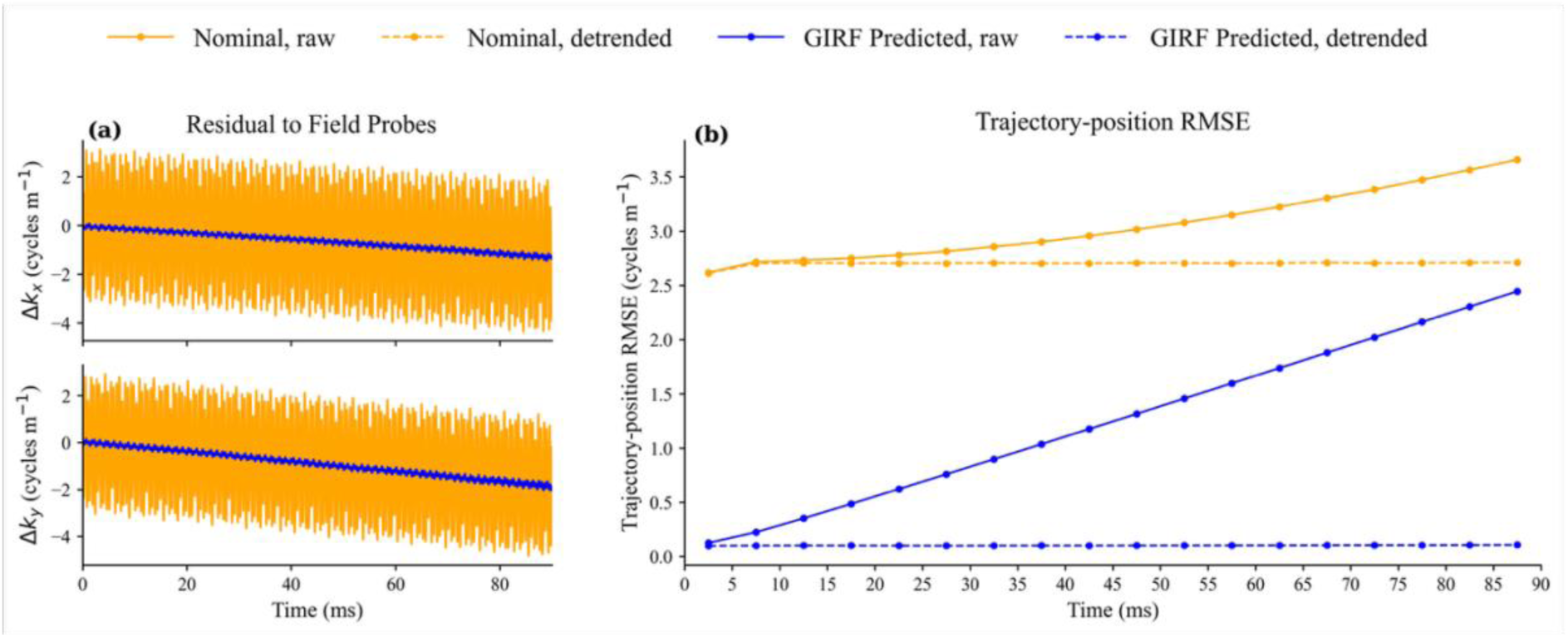
Long-duration trajectory validation over the full 90 ms acquisition window supported by NMR field probes. **(a)** Residual trajectory error relative to the field probes measurement for the nominal (orange) and GIRF-predicted (blue) trajectories, demonstrating a common linear trend over the extended acquisition. **(b)** RMS trajectory position error calculated over consecutive 5 ms intervals before (solid lines) and after (dashed lines) linear detrending.

To investigate the contribution of this common linear trend, the residual trajectories were linearly detrended prior to recalculating the RMS trajectory error. The detrended RMS was then evaluated over consecutive 5 ms intervals throughout the readout (Figure 5b). Following linear detrending, the RMS trajectory error of both the nominal and GIRF-predicted trajectories remained approximately constant throughout the acquisition, with the GIRF prediction exhibiting a substantially lower RMS error (0.10 cycles m^-1^) than the nominal trajectory (2.71 cycles m^-1^). These results suggest that the increasing RMS trajectory error observed over the full acquisition window is dominated by a slowly varying component present in the measured trajectory but absent from both the nominal and GIRF-predicted trajectories. The approximately constant detrended RMS error indicates that the measured GIRFs continue to accurately predict the dynamic component of the trajectory throughout the acquisition.

Figure 6 compares the B_0_ phase measured using the Skope system with that predicted from the measured GIRF. The raw field probes data exhibits a strong linear trend corresponding to a constant frequency offset of 10.1 Hz. As phase is the time integral of frequency, a constant frequency offset produces a linearly increasing phase over time. This offset most likely arose from a slight difference between the frequency reference used by the Skope system during trajectory monitoring and the scanner centre frequency used during initial calibration of the field probes, rather than from the gradient response itself. The linear trend was therefore removed prior to comparison. Following detrending, the measured and GIRF-predicted B_0_ phase showed strong agreement over the full 90 ms readout, with an RMS error of 0.052 radians.

**Figure 6:**
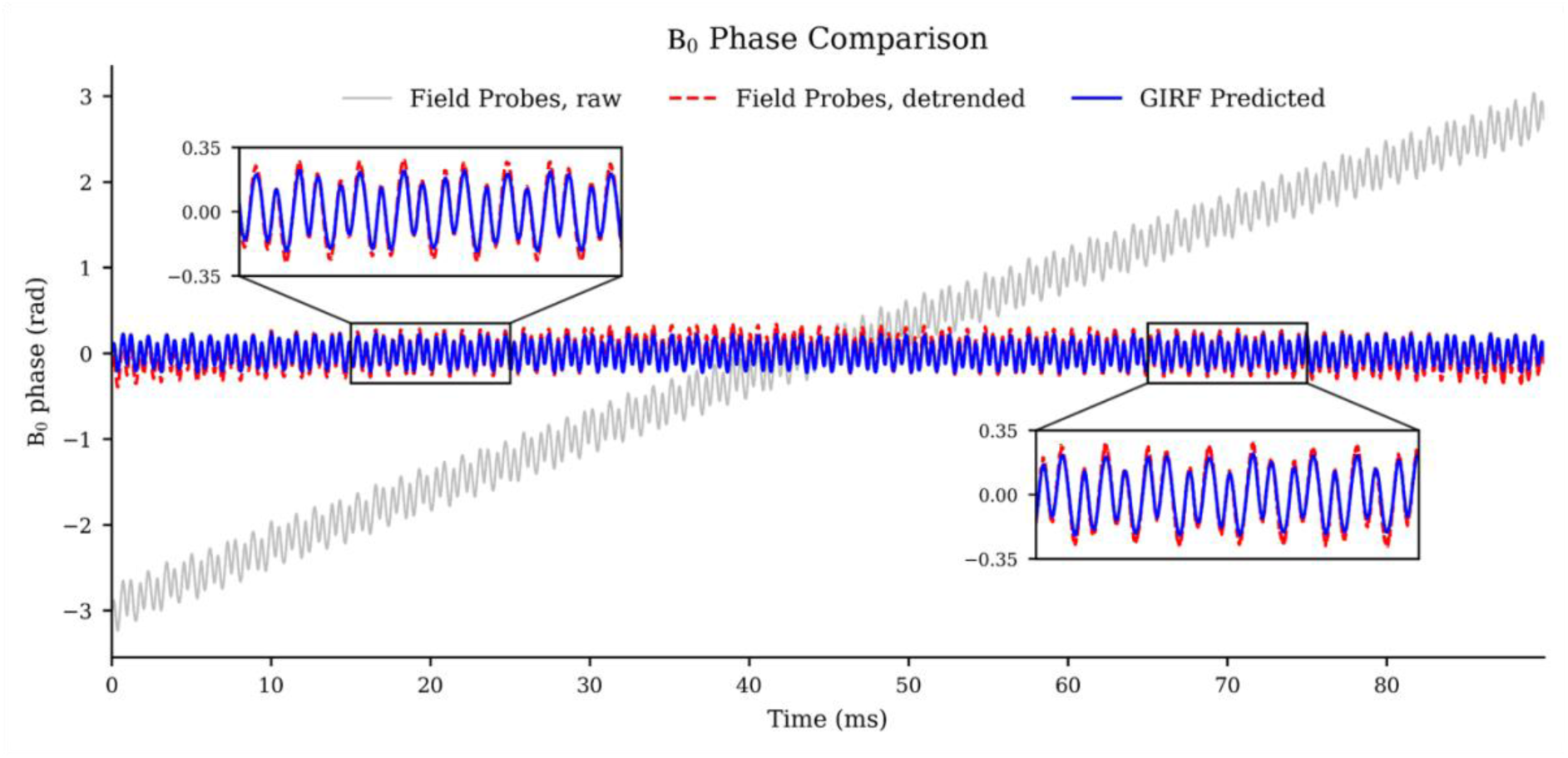
Comparison of the B_0_ phase measured using the NMR field probes and predicted from the measured GIRF for a rosette trajectory in the axial plane. The raw field probes phase (grey) exhibits a linear trend arising from a constant frequency offset between the field probes reference and the scanner centre frequency. After removal of the linear trend (red), strong agreement is observed with the GIRF-predicted B_0_ phase (blue) over the full 90 ms readout. For visualisation, the raw field probes data has been plotted with zero mean.

## Discussion

This work presents an open-source, end-to-end framework for phantom-based measurement of the GIRF. The framework integrates vendor-independent pulse sequence generation, automated data processing, and GIRF estimation into a reproducible workflow requiring only standard MRI hardware and a spherical water phantom. Validation against gold-standard, independent field probe measurements demonstrated that the estimated GIRF accurately predicts non-Cartesian k-space trajectories for MRSI acquisitions employing high-duty-cycle gradient waveforms and extended readout durations.

The principal advantage of the proposed framework is that it provides an openly available, reproducible workflow for GIRF measurement, without requiring specialised field monitoring hardware. By integrating sequence generation, data processing, and GIRF estimation into a single framework, it reduces the need for laboratory-specific software development and promotes reproducible implementation. The phantom-based measurement approach also enables longer readout durations than are conventionally achievable using an NMR field camera, providing improved low-frequency spectral resolution that is primarily limited by the (controllable) phantom T2* and the total acquisition duration. This increased spectral resolution is particularly beneficial for applications requiring accurate characterisation of the low-frequency gradient system behaviour, such as MR spectroscopy.

These advantages are accompanied by several practical trade-offs. First, estimation of higher-order spherical harmonic components using the default triangular-blip protocol requires relatively long data collection times, particularly with 2D phase encoding. In practice, however, GIRF characterisation is typically performed as a one-time scanner calibration procedure. Previous studies have demonstrated good long-term stability of scanner-specific GIRFs over periods of several months to years [9,17], suggesting that routine remeasurement is generally unnecessary except following scanner modifications. Secondly, although the current implementation employs triangular blips, the framework has been designed such that alternative excitation waveforms, such as chirp-based acquisitions [22], could be incorporated without requiring modification to the downstream processing pipeline. Finally, the phantom-based measurement approach exhibits low sensitivity to the weaker higher-order (second-order and higher) field components. In practice, reliable estimation is therefore limited to the zeroth- and first-order spherical harmonic components, reflecting both the reduced SNR of the higher-order terms and the intrinsic sensitivity of the phantom-based measurement approach.

Although field probes were used solely for independent validation in the present work, field monitoring hardware can also be used for rapid GIRF measurements using dedicated blip- or chirp-based acquisition schemes [29,30]. GIRF measurements can be completed in approximately one minute which better supports estimation of higher-order spherical harmonic components. The framework presented here is intended as a complementary approach, providing an openly available and reproducible alternative using only standard MRI hardware, while also permitting longer acquisition windows for applications where improved low-frequency spectral resolution is advantageous.

Validation against independent field probe measurements demonstrated that the proposed framework accurately estimates scanner-specific GIRFs suitable for trajectory prediction. Across all trajectories and imaging orientations, the GIRF-predicted trajectories consistently exhibited substantially lower RMS trajectory error than the corresponding nominal trajectories. Strong agreement was also observed between the measured and GIRF-predicted B_0_ phase following removal of a linear phase trend arising from a small frequency offset between the field probe reference and the scanner centre frequency. Together, these results demonstrate that the proposed phantom-based framework accurately estimates scanner-specific GIRFs and their associated field response for trajectory prediction.

Over longer acquisition windows, a linear trend became apparent in the residual trajectory error. This trend originates from a slowly varying component present in the field probe measurements that is not reproduced by either the nominal or GIRF-predicted trajectories. The approximately constant detrended RMS trajectory error demonstrates that the dynamic component of the measured trajectory continues to be accurately predicted throughout the acquisition, indicating that the observed drift does not arise from inaccuracies in the measured GIRF itself. Instead, the remaining discrepancy is likely associated with behaviour that is not represented by the linear time-invariant assumptions underlying the GIRF model. Possible contributors include non-linear gradient amplifier behaviour [11,31], including amplifier saturation during the prolonged high-duty-cycle readouts used in this study, together with other slowly varying effects such as thermal drift. These effects are directly captured by field monitoring hardware but fall outside the assumptions of the conventional linear time-invariant GIRF-based trajectory prediction.

Previous work has similarly identified circumstances in which the conventional linear time-invariant GIRF model may not fully describe the gradient system response. For example, thermal loading has been shown to induce measurable changes in the GIRF, reducing the validity of the time-invariant assumption [28]. Importantly, these effects do not necessarily require repeated GIRF measurements during routine operation, but instead motivate continued refinement of GIRF-based trajectory prediction models, including thermally informed extensions. Nevertheless, these limitations are inherent to the underlying prediction model rather than to the phantom-based measurement framework presented here.

## Conclusion

We have presented an open-source framework for end-to-end phantom-based measurement of the Gradient Impulse Response Function (GIRF). The framework combines vendor-independent pulse sequence generation, phantom-based data acquisition, automated data processing, and GIRF estimation into a reproducible workflow requiring only standard MRI hardware. Validation against independent field probe measurements demonstrated accurate prediction of non-Cartesian trajectories, confirming that the framework accurately captures the dynamic response of the gradient system. By making phantom-based GIRF measurement openly accessible and reproducible, this work lowers the barrier to routine scanner-specific GIRF characterisation and facilitates broader adoption of GIRF-based methods across the MRI community.

## Data and Code Availability Statement

The data and code used in this work are made available online. Code relating to the open-source framework is available at https://github.com/jbbacon/GIRF_PE_Python (version: Release 1.1.0, specific git hash 5eb8a28) with data and a permanent record available at https://doi.org/10.5281/zenodo.15350583. The GitHub repository will be actively maintained and supported for a minimum of five years following publication, including bug fixes, compatibility updates where practical, and documentation improvements.

## Acknowledgments

This research was funded by the Wellcome grant (225924/Z/22/Z) and supported by the NIHR Oxford Health Biomedical Research Centre (NIHR203316). The views expressed are those of the authors and not necessarily those of the NIHR or the Department of Health and Social Care. The Centre for Integrative Neuroimaging was supported by core funding from the Wellcome Trust (203139/Z/16/Z and 203139/A/16/Z). J.B.B. and P.J. acknowledge support from the Vivensa Foundation. P.J. also acknowledges support from the NIHR Oxford Biomedical Research Centre (NIHR203311). For the purpose of open access, the author has applied a CC BY public copyright license to any Author Accepted Manuscript version arising from this submission. The authors would like to thank Mara Cercignani for her assistance with project coordination and logistics.

## Figure Captions

**Figure 7:** Overview of the proposed end-to-end framework for phantom-based GIRF measurement. A vendor-independent sequence is generated using PyPulseq *[21]* and used to acquire thin-slice, phase-encoded measurements from a spherical phantom. The acquired data are processed through an automated reconstruction pipeline to estimate the output field responses, which are subsequently transformed into the frequency domain to calculate the GIRF. The workflow is divided into three principal stages: pulse sequence generation (grey), automated data processing (light green), and GIRF estimation (blue).

**Figure 8:** Thin-slice acquisition and voxel selection used for phantom-based GIRF measurement. Left: Four offset, phase-encoded thin slices provide spatial sampling of the gradient-induced field response throughout the spherical phantom. Right: Voxels are selected using an intensity threshold prior to spherical harmonic fitting. The threshold preferentially excludes the low-signal voxels near the phantom boundary. Figure adapted from Rahmer et al. *[18]*

**Figure 9:** Magnitude of the GIRFs estimated on a 3T Connectom. **(a-c)** Full frequency response of the zeroth- and first-order spherical harmonic components for input gradient axes *j* = *x*, *y* and *z*, respectively. **(d-i)** Enlarged views of the corresponding self-response **(d-f)** and cross-response/*B*_0_ terms **(g-i)**, each displayed between 0 and 1500 Hz. All GIRFs were estimated using a reconstructed readout duration of 70 ms. Output field spherical harmonic components (*m*) are colour coded as *B*_0_ (red), *x* (blue), *y* (orange), and *z* (green).

**Figure 10:** Comparison of the nominal, GIRF-predicted and field probes-measured rosette trajectories acquired in the axial plane. Only the first 6.6 ms of the readout is displayed. **(a,b)** *k_x_* and *k_y_* trajectories as a function of time, with enlarged views highlighting the agreement between the GIRF-predicted and measured trajectories. **(c,d)** Residual trajectory error relative to the field probes measurement for the nominal and GIRF-predicted trajectories. **(e)** Two dimensional *k_x_*-*k_y_* trajectory. **(f)** Enlarged view around the central region of the trajectory. Trajectories are colour coded as Nominal (orange), GIRF-predicted (blue), and Field Probes measured (red).

**Figure 11:** Long-duration trajectory validation over the full 90 ms acquisition window supported by NMR field probes. **(a)** Residual trajectory error relative to the field probes measurement for the nominal (orange) and GIRF-predicted (blue) trajectories, demonstrating a common linear trend over the extended acquisition. **(b)** RMS trajectory position error calculated over consecutive 5 ms intervals before (solid lines) and after (dashed lines) linear detrending.

**Figure 12:** Comparison of the B_0_ phase measured using the NMR field probes and predicted from the measured GIRF for a rosette trajectory in the axial plane. The raw field probes phase (grey) exhibits a linear trend arising from a constant frequency offset between the field probes reference and the scanner centre frequency. After removal of the linear trend (red), strong agreement is observed with the GIRF-predicted B_0_ phase (blue) over the full 90 ms readout. For visualisation, the raw field probes data has been plotted with zero mean.

Supporting Figure S1: Phase of the self-term of the GIRFs measured on a 3T Connectom. **(a-c)** Full frequency response for the input gradient axis *j* = *x*, *y* and *z*, respectively. **(d-f)** Enlarged view of the phase response between -1500 and 1500 Hz.

Supporting Figure S2: Comparison of the nominal, GIRF-predicted and field probes-measured rosette trajectories acquired in the coronal and sagittal planes. Only the first 6.6 ms of the readout is displayed.

Supporting Figure S3: Comparison of the nominal, GIRF-predicted and field probes-measured spiral trajectories acquired in the axial, coronal and sagittal planes. Only the first 7.8 ms of the readout is displayed.

