## Supporting Information for "Open-Source Framework for Measurement of the Gradient Impulse Response Function"

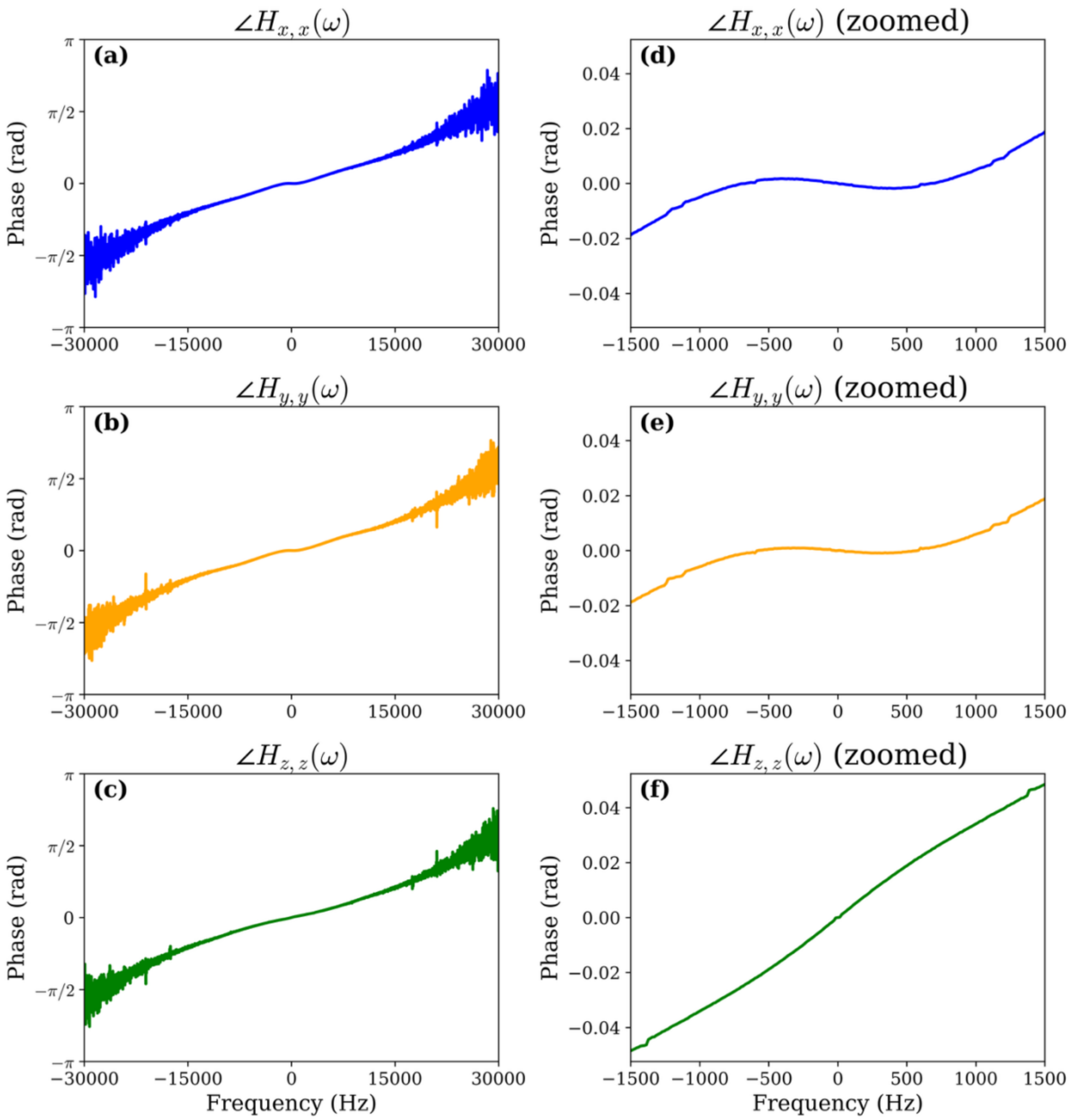


Supporting Figure S1: Phase of the self-term of the GIRFs measured on a 3T Connectom **(a-c)** Full frequency response for the input gradient axis $j = x, y$ and $z$, respectively. **(d-f)** Enlarged view of the phase response between -1500 and 1500 Hz.

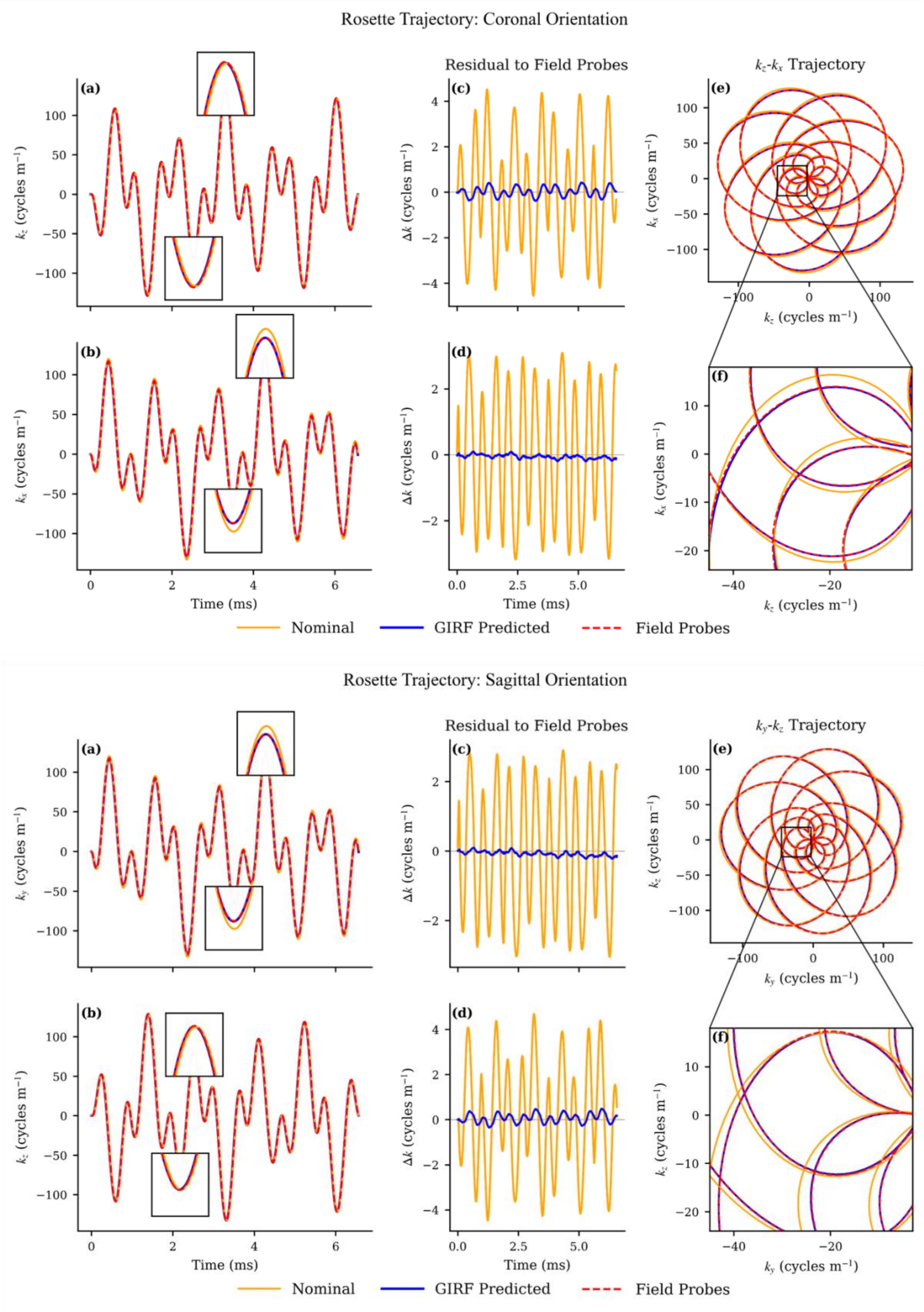


Supporting Figure S2: Comparison of the nominal, GIRF-predicted and field probes-measured rosette trajectories acquired in the coronal and sagittal planes. Only the first 6.6 ms of the readout is displayed.

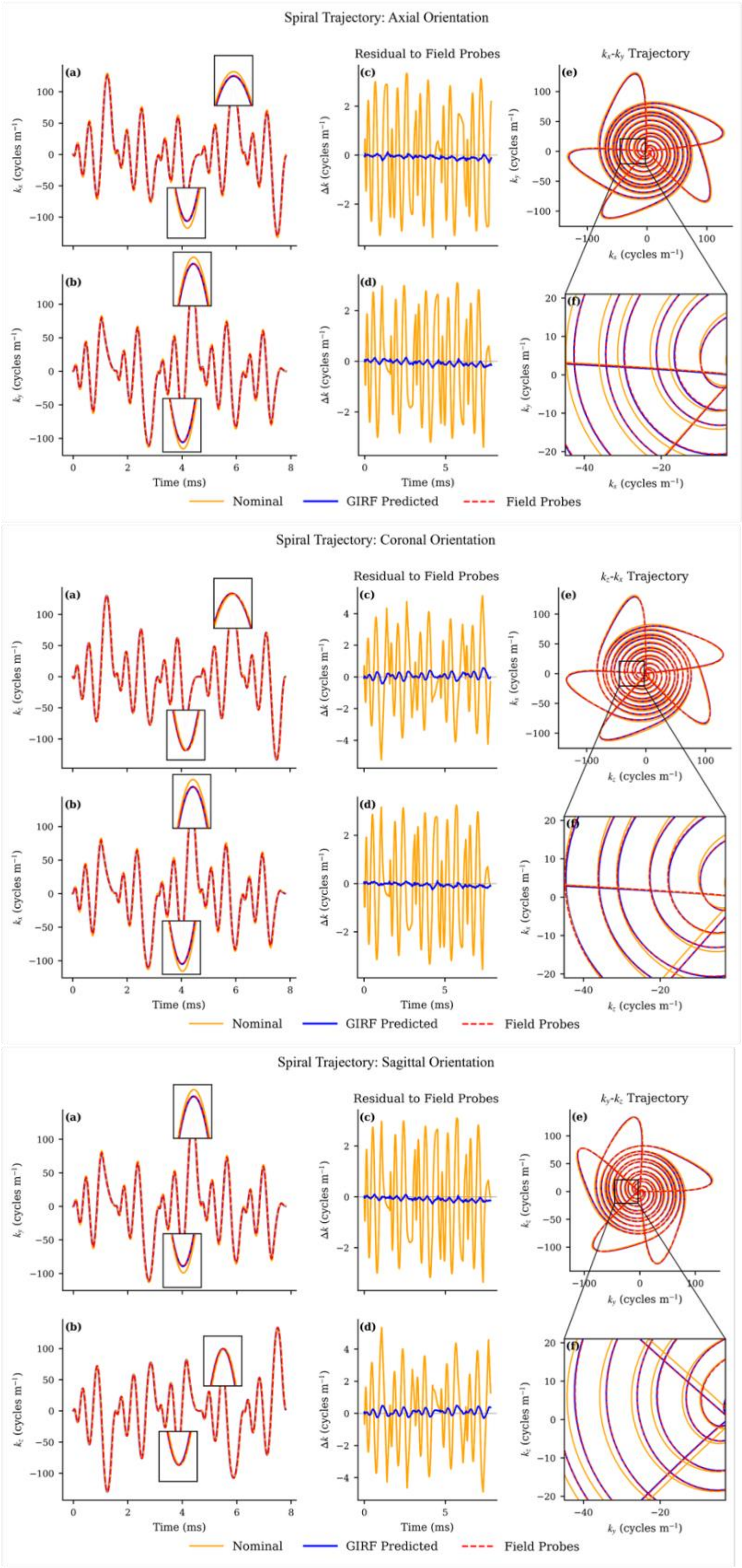


Supporting Figure S3: Comparison of the nominal, GIRF-predicted and field probes-measured spiral trajectories acquired in the axial, coronal and sagittal planes. Only the first 7.8 ms of the readout is displayed.
